# Evolved increases in hemoglobin-oxygen affinity and Bohr effect coincided with the aquatic specialization of penguins

**DOI:** 10.1101/2020.11.17.387597

**Authors:** Anthony V. Signore, Michael S. Tift, Federico G. Hoffmann, Todd. L. Schmitt, Hideaki Moriyama, Jay F. Storz

**Affiliations:** School of Biological Sciences, University of Nebraska, Lincoln, NE 68588, USA; Department of Biology and Marine Biology, University of North Carolina, Wilmington, NC 28403, USA; Department of Biochemistry, Molecular Biology, Entomology, and Plant Pathology, Mississippi State University, MS 39762, USA; Veterinary Services, SeaWorld of California, San Diego, CA 92109, USA

**Keywords:** Hemoglobin, hypoxia, diving, penguins, Bohr effect, adaptation

## Abstract

Dive capacities of air-breathing vertebrates are dictated by onboard O_2_ stores, suggesting that physiological specializations of diving birds like penguins may have involved adaptive changes in convective O_2_ transport. It has been hypothesized that increased hemoglobin (Hb)-O_2_ affinity improves pulmonary O_2_ extraction and enhance capacities for breath-hold diving. To investigate evolved changes in Hb function associated with the aquatic specialization of penguins, we integrated comparative measurements of whole-blood and purified native Hbs with protein engineering experiments based on site-directed mutagenesis. We reconstructed and resurrected ancestral Hbs representing the common ancestor of penguins and the more ancient ancestor shared by penguins and their closest nondiving relatives (order Procellariiformes, which includes albatrosses, shearwaters, petrels, and storm petrels). These two ancestors bracket the phylogenetic interval in which penguin-specific changes in Hb function would have evolved. The experiments revealed that penguins evolved a derived increase in Hb-O_2_ affinity and a greatly augmented Bohr effect (reduced Hb-O_2_ affinity at low pH). Although an increased Hb-O_2_ affinity reduces the gradient for O_2_ diffusion from systemic capillaries to metabolizing cells, this can be compensated by a concomitant enhancement of the Bohr effect, thereby promoting O_2_ unloading in acidified tissues. We suggest that the evolved increase in Hb-O_2_ affinity in combination with the augmented Bohr effect maximizes both O_2_ extraction from the lungs and O_2_ unloading from the blood, allowing penguins to fully utilize their onboard O_2_ stores and maximize underwater foraging time.

## Introduction

In air-breathing vertebrates, diving capacities are dictated by onboard O_2_ stores and the efficiency of O_2_ use in metabolizing tissues (1). In fully aquatic taxa, selection to prolong breath-hold submergence and underwater foraging time may have promoted adaptive changes in multiple components of the O_2_-transport pathway, including oxygenation properties of hemoglobin (Hb). Vertebrate Hb is a tetrameric protein that is responsible for circulatory O_2_ transport, loading O_2_ at pulmonary capillaries and unloading O_2_ in the systemic circulation via quaternary structural shifts between a high affinity (predominately oxygenated) relaxed (R-) state and a low affinity (predominately deoxygenated) tense (T-) state (2). While this mechanism of respiratory gas transport is conserved in all vertebrate Hbs, amino acid variation in the constituent α- and β-type subunits may alter intrinsic O_2_ affinity and the responsiveness to changes in temperature, red cell pH, and red cell concentrations of allosteric cofactors (non-heme ligands that modulate Hb-O_2_ affinity by preferentially binding and stabilizing the deoxy T conformation) (3, 4).

While the quantity of Hb is typically increased in the blood of diving birds and mammals in comparison with their terrestrial relatives, there is no consensus on whether evolved changes in Hb–O_2_ affinity have contributed to enhanced diving capacities (1). It has been hypothesized that increased Hb-O_2_ affinity may improve pulmonary O_2_ extraction in diving mammals, thereby enhancing diving capacity (5), but more comparative data are needed to assess evidence for an adaptive trend (6, 7). Experimental measurements on whole-blood suggest that the emperor penguin (*Aptenodytes forsteri*) may have a higher blood-O_2_ affinity relative to nondiving waterbirds, a finding that has fostered the view that this is a property that characterizes penguins as a group (8–10). However, blood-O_2_ affinity is a highly plastic trait that is influenced by changes in red cell metabolism and acid-base balance, so measurements on purified Hb are needed to assess whether observed species differences in blood-O_2_ affinity stem from genetically based changes in the oxygenation properties of Hb. Moreover, even if species differences in Hb-O_2_ affinity are genetically based, comparative data from extant taxa do not reveal whether observed differences are attributable to a derived increase in penguins, a derived reduction in their nondiving relatives, or a combination of changes in both directions.

To investigate evolved changes in Hb function associated with the aquatic specialization of penguins, we integrated experimental measurements of whole-blood and purified native Hbs with evolutionary analyses of globin sequence variation. To characterize the mechanistic basis of evolved changes in Hb function in the stem lineage of penguins, we performed protein engineering experiments on reconstructed and resurrected ancestral Hbs representing (*i*) the common ancestor of penguins and (*ii*) the more ancient ancestor shared by penguins and their closest nondiving relatives (order Procellariiformes, which includes albatrosses, shearwaters, petrels, and storm petrels) (Figure 1). These two ancestors bracket the phylogenetic interval in which penguin-specific changes in Hb function would have evolved.

**Figure 1.**
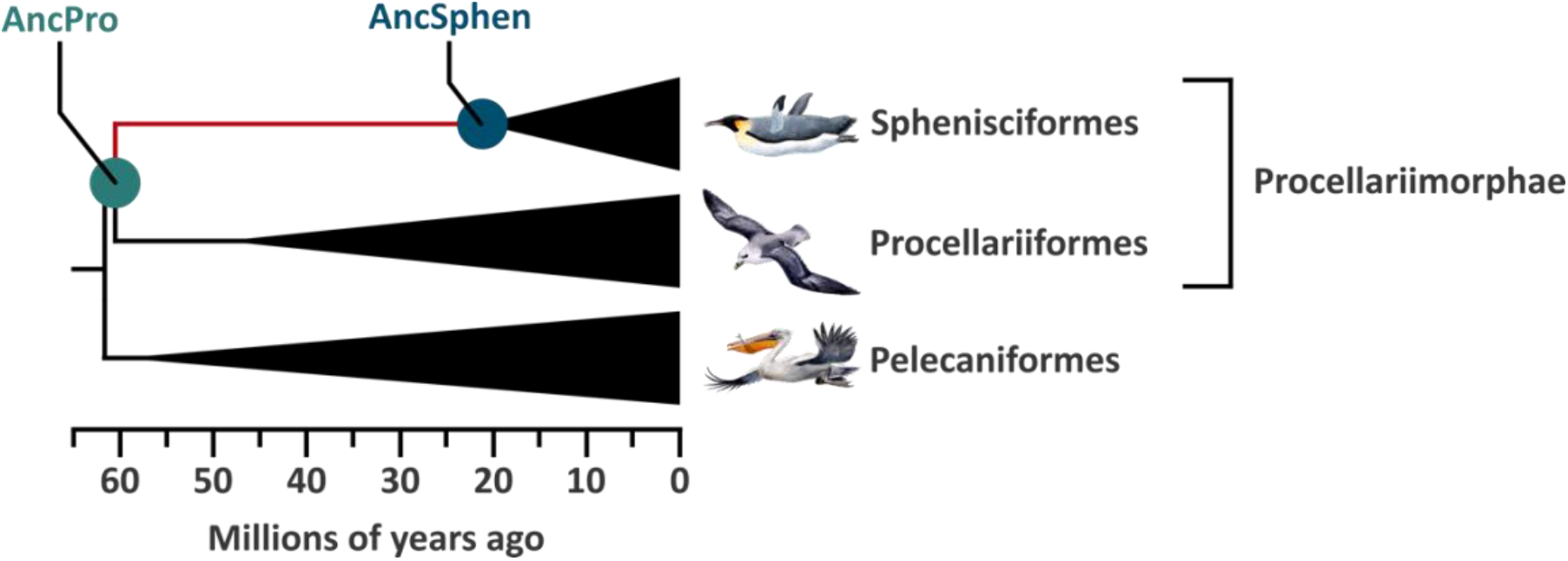
Diagrammatic phylogeny showing the relationship between Sphenisciformes, Procellariifomes and Pelecaniformes. Ancestral hemoglobins were reconstructed for the two indicated nodes: the common ancestor of Sphenisciformes (AncSphen) and the common ancestor of Procellariimorphae (AncPro), the super order that contains Sphenisciformes and Procellariifomrmes. Divergence times are adapted from Claramunt and Cracraft (51).

## Results and Discussion

### Oxygen binding properties of penguin whole-blood and purified Hbs

Using blood samples from multiple individuals of six penguin species, we measured the partial pressure of O_2_ (*P*_O2_) at 50% saturation (*P*_50_) for whole-blood and purified Hbs in the absence (stripped) and presence of allosteric cofactors (+KCl +IHP [inositol hexaphosphate]) (Figure 2). Whole-blood *P*_50_ values were similar across all penguins, averaging 33.3±1.1 torr (Figure 2; Table S1), consistent with previously published data for emperor, Adélie, chinstrap, and gentoo penguins (8, 9, 11). Similarly, measured O_2_-affinities for purified Hbs exhibited very little variation among species, both in the presence and absence of allosteric cofactors (Figure 2; Table S1). Penguins express a single Hb isoform during postnatal life (HbA), in contrast to the majority of other bird species that express one major and one minor isoform (HbA and HbD, respectively) (12, 13). The lack of variation in Hb-O_2_ affinity among penguins is consistent with the low level of amino acid variation in the α- and β-chains (Figure S1). The experiments revealed that penguin Hbs exhibit a remarkably large shift in the magnitude of the Bohr effect (i.e. the reduction in Hb–O_2_ affinity in response to reduced pH) with the addition of allosteric cofactors (Table S1). The average Bohr effect of penguin Hb more than doubles with the addition of allosteric cofactors, from −0.21±0.03 to −0.53±0.04 (Table S1).

**Figure 2.**
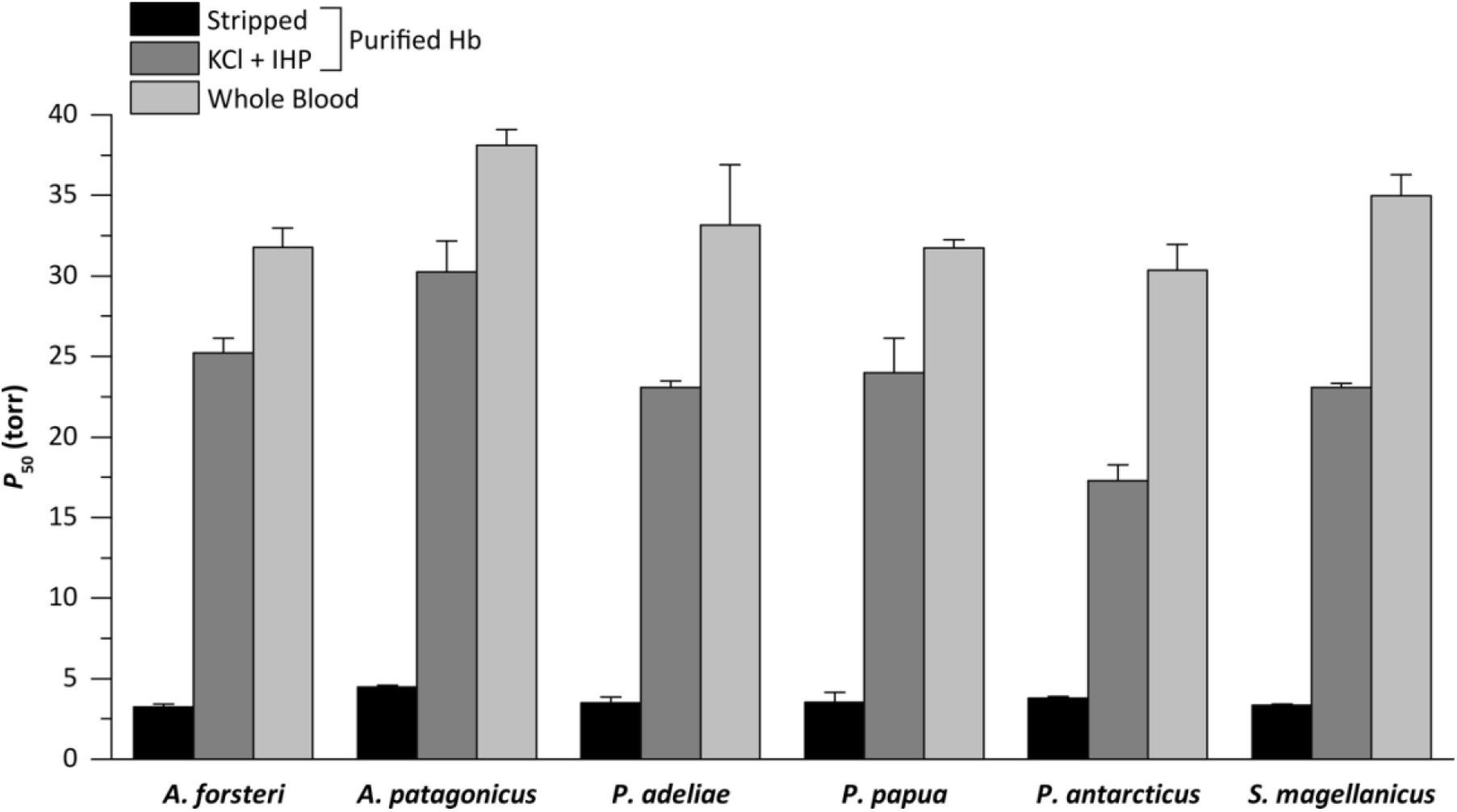
Oxygen tensions at half saturation (*P*_50_) for penguin whole-blood and purified hemoglobins at 37°C, in the absence (Stripped) and presence of 100 mM KCl and 0.2 mM inositol hexaphosphate (+KCl +IHP). The higher the *P*_50_, the lower the Hb-O_2_ affinity. Whole-blood *P*_50_ values are presented as mean±S.E (n=3). Purified Hb *P*_50_ values are derived from plots of log*P*_50_ vs. pH, where a linear regression was fit to estimate *P*_50_ at exactly pH 7.40 (± S.E. of the regression estimate).

Our experimental results indicate that penguins have a generally higher Hb-O_2_ affinity than other birds (12, 14–22), consistent with previous suggestions based on measurements on whole-blood (8, 9, 23–25). Whole-blood O_2_-affinities of the six examined penguin species (30.4 to 38.1 torr at 37°C, pH 7.40) were uniformly higher than that from a representative member of Procellariformes, the southern giant petrel (*Macronectes giganteus;* 42.5 torr at 38°C, pH 7.40) (9). Similarly, numerous high-altitude bird species have convergently evolved increased Hb–O_2_ affinities (17, 18, 20, 21), which appears to be adaptive because it helps safeguard arterial O_2_ saturation in spite of the reduced *P*_O2_ of inspired air (26–28). The difference in blood *P*_50_’s between penguins and the southern giant petrel is generally much greater in magnitude than differences in Hb *P*_50_ between closely related species of low- and high-altitude birds (17, 18, 20, 21). Similar to the case of other diving vertebrates (29), the Bohr effect of penguin Hb also greatly exceeds typical avian values.

### Ancestral protein resurrection

In principle, the observed difference in Hb-O_2_ affinity between penguins and their closest non-diving relatives could be explained by a derived increase in Hb-O_2_ affinity in the penguin lineage (the generally assumed adaptative scenario), a derived reduction in the stem lineage of Procellariformes (the nondiving sister group), or a combination of changes in both directions. To test these alternative hypotheses, we reconstructed the Hbs of the common ancestor of penguins (AncSphen) and the more ancient common ancestor of Procellariimorphae (the superorder comprising Sphenisciformes [penguins] and Procellariiformes; AncPro) (Figures 1, S2, S3 and S4). We then recombinantly expressed and purified the ancestral Hbs to perform *in vitro* functional tests. Measurements of O_2_-equilibrium curves revealed that the AncSphen Hb has a significantly higher O_2_-affinity than that of AncPro (Figure 3), indicating that penguins evolved a derived increase in Hb-O_2_ affinity. In the presence of allosteric cofactors, the *P*_50_ of AncSphen is much lower (O_2_-affinity is higher) compared to AncPro (11.8 vs. 20.2 torr). Much like the evolved increases in Hb-O_2_ affinity in high-altitude birds (18, 20–22), the increased O_2_-affinity of penguin Hb is attributable to an increase in intrinsic affinity rather than a reduced responsiveness to allosteric cofactors, as the Hb–O_2_ affinity difference between AncSphen and AncPro persists in the presence and absence of Cl^−^ and IHP (Figure 3).

**Figure 3.**
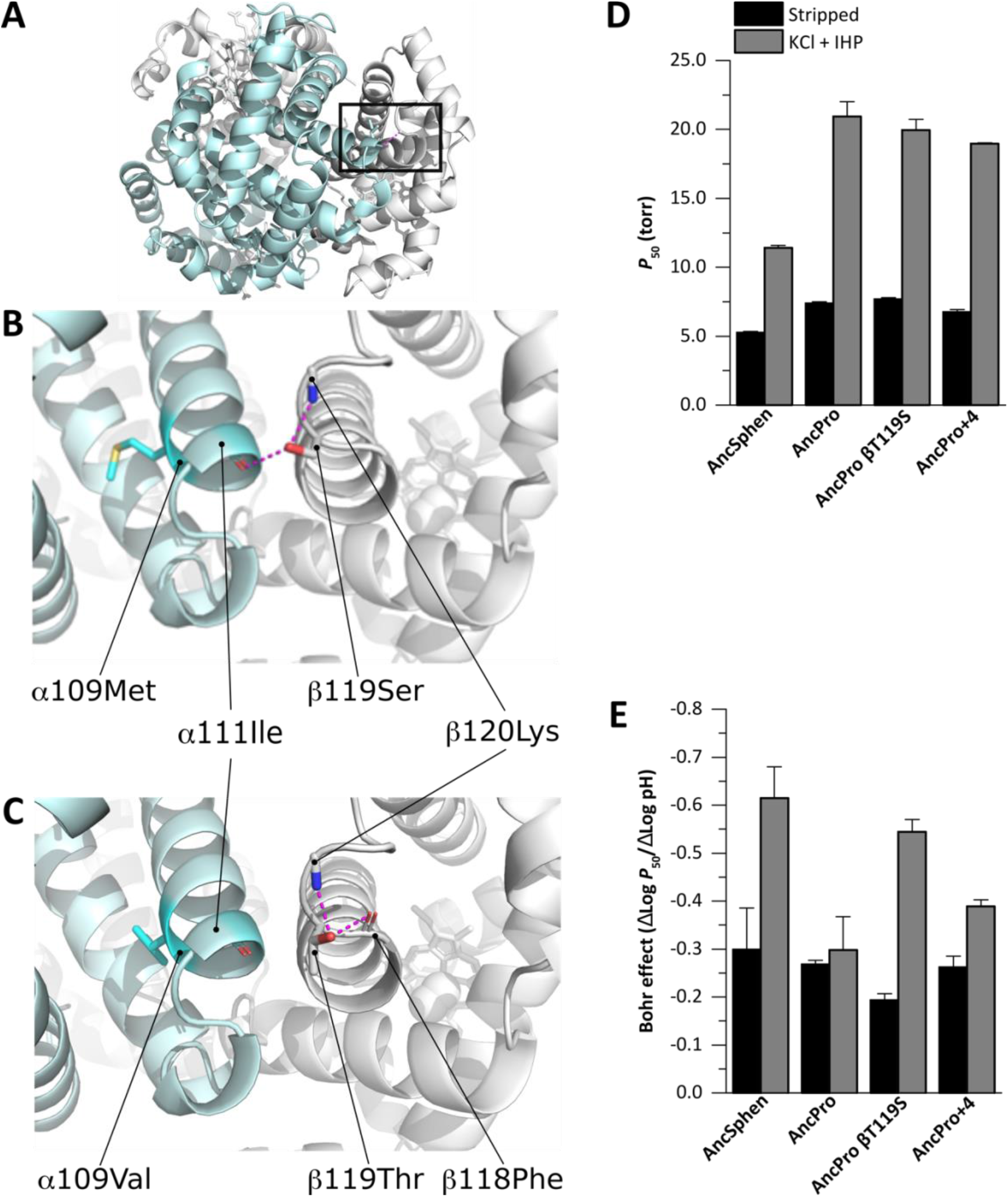
Structural (A–C) and physiological effects (D, E) of amino acid substitutions in the reconstructed Hb proteins of the penguin ancestor (AncSphen) and the last common ancestor penguins shared with Procellariiformes (AncPro). A) Molecular model of the AncSphen Hb tetramer where the black box indicates the regions highlighted in panels B and C. B) Molecular model of AncSphen Hb showing inter-subunit stabilizing H-bonds (pink) between β119Ser and both α111Ile and β120Lys. C) Molecular model of AncPro Hb showing that replacement of β119Ser with Thr removes the inter-subunit stabilizing H-bonds. D) Hb-O_2_ affinity (as measured by *P*_50_, the O_2_ tension at half saturation) of AncSphen, AncPro, and two mutant rHbs with penguin-specific amino acid replacements introduced on the AncPro background (AncProβ119Ser and AncPro+4). See text for explanation regarding the choice of candidate sites for mutagenesis experiments. Measurements were performed on Hb solutions (0.1 mM Hb in 0.1 M HEPES/0.5 mM EDTA) at 37°C in the absence (stripped) and presence of 0.1 M KCl and 0.2 mM inositol hexaphosphate (+KCl +IHP). *P*_50_ values are derived from plots of log*P*_50_ vs. pH, where a linear regression was fit to estimate *P*_50_ at exactly pH 7.40 (± S.E. of the regression estimate). E) Bohr coefficients (ΔLog P50/ΔLog pH) were estimated from plots of log*P*_50_ vs. pH, where the Bohr effect is represented by the slope of a linear regression (± S.E. of the slope estimate).

In addition to the derived increase in Hb–O_2_ affinity, comparisons between AncSphen and AncPro also revealed that the Hb of penguins evolved an enhanced responsiveness to pH (Bohr effect). Under stripped conditions, the Bohr effect of AncSphen and AncPro (−0.30±0.09 and −0.27±0.1, respectively) were highly similar to one another and were similar to values measured for native penguin Hbs under the same conditions (Figure 3E; Table S1). However, in the presence of allosteric cofactors the Bohr effect of AncSphen increases more than two-fold (similar to that of native penguin Hbs), whereas that of AncPro shows little change (Figure 3E), demonstrating that penguins evolved an increased cofactor-linked Bohr effect following divergence from their non-diving relatives. An increased Hb-O_2_ affinity is expected to reduce the gradient for O_2_ diffusion from systemic capillaries to the cells of metabolizing tissues, and an increased Bohr effect can compensate for this by reducing Hb-O_2_ affinity at low pH, thereby promoting O_2_ unloading in acidified tissues. A similar augmentation of the Bohr effect was recently documented in the Hb of high-altitude Tibetan canids (30). In summary, the Hbs of penguins evolved an increase in O_2_-affinity and enhanced Bohr effect in association with other physiological and morphological specializations for a more fully aquatic existence.

### Tests of positive selection

Given that joint increases in the O_2_-affinity and Bohr effect of penguin Hb represent derived character states, we performed a molecular evolution analysis to test for evidence of positive selection in the α- and β-globin genes. Specifically, we tested for an accelerated rate of amino acid substitution in the stem lineage of penguins (the branch connecting Anc Procellariimorphae [AncPro] to the common ancestor of penguins [AncSphen]) using the branch-sites test. This test revealed no evidence for an accelerated rate of amino acid substitution in the stem lineage of penguins (Table S2), and a clade test revealed no significant variation in substitution rate among different penguin lineages (Table S3). Thus, if the increased Hb-O_2_ affinity of penguins represents an adaptation that evolved via positive selection, the nature of the causative changes did not produce a detectable statistical signature in the α- and β-type globin genes.

### Molecular modelling

We used molecular modelling to identify which specific amino acid substitutions may be responsible for the increased Hb-O_2_ affinity of AncSphen relative to AncPro. Of the 17 amino acid substitutions that distinguish AncSphen and AncPro, our analyses identified four substitutions that could potentially alter O_2_-binding properties. The substitution Thrβ119Ser in the branch leading to AncSphen affects the stabilization of R-state (oxygenated) Hb. Specifically, the hydroxyl group of β119Ser in helix G is oriented toward the subunit interface by forming a hydrogen-bond with β120Lys, which permits an intersubunit contact with α111Ile (Figure 3A,B). This bond between β119Ser and α111Ile stabilizes the R-state conformation by clamping the intersubunit motions, which is predicted to increase Hb-O_2_ affinity by raising the free energy of the oxygenation-linked allosteric R→T transition in quaternary structure. Additionally, our model identified three other amino acid substitutions, αA138S, βA51S and βI55L, that create intersubunit contacts and further stabilize the R-state conformation.

### Testing causative substitutions

To test model-based predictions about the specific substitutions that are responsible for the increased O_2_-affinity of penguin Hb, we used site-directed mutagenesis to introduce combinations of mutations at four candidate sites on the AncPro background. We first tested the effect of a single mutation whereby β119Thr was replaced with Ser (AncProβT119S). We then tested the net effect of mutations at all 4 sites on the AncPro background (AncPro+4; αA138S, βA51S, βI55L, and βT119S). The protein engineering experiments revealed that βT119S produced a negligible individual effect on Hb-O_2_ affinity when introduced on the AncPro background, but it produced an appreciable increase in the Bohr effect (Figure 3D, E). The 4 mutations in combination produced a modest increase in Hb-O_2_ affinity and a more pronounced increase in the Bohr effect, but they did not fully recapitulate observed differences between AncPro and AncSphen in either of these properties (Figure 3). These data suggest the evolved functional changes in penguin Hb must be attributable to the net effect of multiple amino acid substitutions at structurally disparate sites.

### Adaptive significance of increased Hb-O_2_ affinity

The key to extending dive times for aquatic vertebrates is to increase O_2_ carrying capacity while keeping metabolic O_2_ demands as low as possible during breath-hold submergence. Submergence induces intense bradycardia and peripheral vasoconstriction, which conserves finite O_2_ stores for tissues that are intolerant to hypoxia (i.e. the central nervous system and heart) (31). O_2_ stores are typically increased in diving vertebrates via increased blood volume, increased blood Hb concentration, increased myoglobin concentration in skeletal muscle, increased muscle mass and, occasionally, increased diving lung volume (1). As deep diving cetaceans and pinnipeds exhale before submergence, their lungs account for less than 10% of total O_2_ stores (1, 32). This reduction in diving lung volume reduces gaseous N_2_ and O_2_, which presumably limits decompression sickness. Conversely, as penguins inhale at the onset of a dive, their diving lung volume accounts for a much larger percentage of total O_2_ stores (19% and 45% for the emperor and Adélie penguins, respectively) (1, 33). Indeed, in diving emperor penguins, O_2_ extraction from pulmonary stores is continuous during submergence (34, 35). An elevated Hb-O_2_ affinity (such as that found in penguins) can maximize O_2_ extraction from pulmonary stores, as greater blood–O_2_ saturation can be achieved at any given parabronchial *P*_O2_. However, while increased Hb-O_2_ affinity may confer more complete transfer of O_2_ from the lungs to the blood, it can inhibit subsequent O_2_ transfer from the blood to the tissues. Despite this, emperor penguins almost completely deplete their circulatory stores during extended dives, as their end-of-dive venous *P*_O2_ can be as low as 1-6 torr (34). The enhanced Bohr effect of penguin Hb should improve O_2_ delivery to working (acidic) tissues, allowing more complete O_2_ unloading of the blood. We suggest this modification works in tandem with increased Hb-O_2_ affinity to maximize both O_2_ extraction from the lungs and O_2_ unloading from the blood, allowing penguins to fully utilize their onboard O_2_ stores and maximize underwater foraging time.

## Materials and Methods

### Blood collection

We collected blood from 18 individual penguins representing six species: *Aptenodytes forsteri*, *A. patagonicus*, *Pygoscelis adeliae*, *P. papua*, *P. antarcticus*, and *Spheniscus magellanicus* (*n*=3 individuals per species). All birds were sampled during routine health checks at SeaWorld of California (San Diego, California). Blood was collected by venipuncture of the jugular vein using Vacutainer^®^ Safety-Lok™ blood collection set (Becton Dickinson, Franklin Lakes, NJ) with 21 G x ¾” (0.8 × 19 mm) needle attached to a heparin blood collection tube (Becton Dickinson). A subsample of whole-blood (200 μl) was set aside for oxygen equilibrium curves (see below) and the remaining blood was centrifuged at 5000xg for 15 minutes. Plasma, buffy coat, and hematocrit fractions from the centrifuged samples were immediately placed in separate tubes and flash frozen at −80°C for future analyses.

### Sequencing of penguin globin genes

RNA was extracted from ~100 μl of flash frozen erythrocytes using an RNeasy Universal Plus Mini Kit (Qiagen). cDNA was synthesized from freshly prepared RNA using Superscript IV Reverse transcriptase (Invitrogen). Gene specific primers used to amplify the α- and β-type globin transcripts were designed from the 5’ and 3’ flanking regions of all publicly available penguin globin genes. PCR reactions were conducted using 1 ml of cDNA template in 0.2 ml tubes containing 25 μl of reaction mixture (0.5 μl of each dNTP (2.5 mM), 2.5 μl of 10x Reaction Buffer (Invitrogen), 0.75 μl of 50 mM MgCl2, 1.25 μl of each primer (10 pmol/μl), 1 μl of Taq polymerase (Invitrogen) and 16.75 μl of ddH2O), using an Eppendorf Mastercycler® Gradient thermocycler. Following a 5-min denaturation period at 94°C, the desired products were amplified using a cycling profile of 94°C for 30 sec; 53-65°C for 30 sec; 72°C for 45 sec for 30 cycles followed by a final extension period of 5 min at 72°C. Amplified products were run on a 1.5% agarose gel and bands of the correct size were subsequently excised and purified using Zymoclean Gel DNA recovery columns (Zymo Research). Gel-purified PCR products were ligated into pCR™4-TOPO® vectors using a TOPO™ TA Cloning™ Kit and were then transformed into One Shot™ TOP10 Chemically Competent E. coli (Thermo Fisher Scientific). Three to six transformed colonies were cultured in 5 ml of LB medium and plasmids were subsequently purified with a GeneJET Plasmid Miniprep kit (Thermo Fisher Scientific). Purified plasmids were sequenced by Eurofins Genomics.

### Sequence analyses

Genomic sequences containing the complete α- and β-globin gene clusters for the emperor penguin (*A. forsteri*), Adélie penguin (*P. adeliae*), northern fulmar (*Fulmarus glacialis*), band-rumped storm-petrel (*Hydrobates castro*), southern giant petrel (*Macronectes giganteus*), flightless cormorant (*Nannopterum harrisi*), crested ibis (*Nipponia nippon*), and the little egret (*Egretta garzetta*) were obtained from GenBank. The α- and β-globin gene clusters from the remaining 19 extant penguin species were obtained from GigaDB (36). Coding sequences of α- and β-globin genes extracted from these genomic sequences were combined with the newly generated cDNA sequences mentioned above (Figure S2). Sequences were aligned using MUSCLE (37) and were then used to estimate phylogenetic trees. The best fitting codon substitution model and initial tree search were estimated using IQ-TREE with the options -st CODON, -m TESTNEW, -allnni, and -bnni (38, 39). Initial trees were then subjected to 1000 ultrafast bootstrap replicates (40). Bootstrap consensus trees (Figure S3) were used to estimate ancestral globin sequences using IQ-TREE with the option -asr (Figures S2 and S4).

### Selection analyses

We tested for selection in the evolution of the penguins’ α- and β-globin genes in a maximum likelihood framework with the codon-based models implemented in the codeml program from the PAML v4.9 suite (41), using the phylogenetic trees described above (see “Sequence analyses”). We used the branch-site and clade models to examine variation in ω, the ratio of the rate of nonsynonymous substitutions per nonsynonymous site, dN, to the rate of synonymous substitutions per synonymous site, dS. We used branch-site model A (42, 43) to test for positive selection in the branch connecting AncPro to AncSphen (the stem lineage of penguins) (Table S2), and we used clade C model (44) to test for selection in the penguin clade using M2a_rel from Weadick and Chang (45) as the null model (Table S3).

### Molecular modeling

Structural modeling was performed on the SWISS MODEL server (46)using graylag goose hemoglobin in oxy form (PDB 1faw). AncPro and AncSphen Hbs had QMEAN values of –0.61 and –0.65, respectively. The root mean square distance of the main chain between template and model (RMSD) values < 0.09Å were considered usable (47). Structural mining and preparation of graphics were performed using the PyMOL Molecular Graphics System, version 2.3.2 (Schrödinger, LLC, New York, NY, USA). Hydrogen bond listing was performed using a PyMol script list_hb.py (Robert L. Campbell, Biomedical and Molecular Sciences, Queen’s University, Canada). The interface binding energy was calculated by the ePISA server (48).

### Construction of Hb expression vectors

Reconstructed ancestral globins were synthesized by GeneArt Gene Synthesis (Thermo Fisher Scientific) after optimizing the nucleotide sequences in accordance with *E. coli* codon preferences. The synthesized globin gene cassette was cloned into a custom pGM vector system along with the methionine aminopeptidase (MAP) gene, as described previously (49). We engineered the Thrβ119Ser substitution by whole plasmid amplification using mutagenic primers and Phusion High-Fidelity DNA Polymerase (New England BioLabs), phosphorylation with T4 Polynucleotide Kinase (New England BioLabs), and circularization with an NEB Quick Ligation Kit (New England BioLabs). All site-directed mutagenesis steps were performed using the manufacture’s recommended protocol. Each plasmid was verified with DNA sequencing by Eurofins genomics.

### Expression and purification of recombinant Hbs

Recombinant Hb expression was carried out in the *E. coli* JM109 (DE3) strain as described previously (15, 49, 50). Bacterial cell lysates were loaded onto a HiTrap SP HP anion exchange column (GE Healthcare) and were then equilibrated with 50 mM HEPES/0.5 mM EDTA (pH 7.0) and eluted with a linear gradient of 0 - 0.25 M NaCl. Hb-containing fractions were then loaded on to a HiTrap Q HP cation exchange column (GE Healthcare) equilibrated with 20 mM Tris-HCl/0.5mM EDTA (pH 8.6) and eluted with a linear pH gradient 0 - 0.25 M NaCl. Eluted Hb factions were concentrated using Amicon Ultra-4 Centrifugal Filter Units (Millipore).

### Sample preparation for O_2_-equilibrium curves

Fresh whole-blood was diluted 1:15 with each individual’s own plasma and oxygen-equilibrium curves were measured immediately after sampling. To obtain stripped hemolysate, 100μl centrifuged red blood cells were added to a 5x volume of 0.01 M HEPES/0.5 mM EDTA buffer (pH 7.4) and incubated on ice for 30 min to lyse the red blood cells. NaCl was added to a final concentration of 0.2 M and samples were centrifuged at 20,000 x g for 10 min to remove cell debris. Hemolysate supernatants and purified recombinant hemoglobins were similarly desalted by passing through a PD-10 desalting column (GE Healthcare) equilibrated with 25 ml of 0.01 M HEPES/0.5mM EDTA (pH 7.4). Eluates were concentrated using Amicon Ultra-4 Centrifugal Filter Units (Millipore). From these concentrated samples, Hb solutions (0.1 mM hemoglobin in 0.1 M HEPES/0.05 M EDTA buffer) were prepared in the absence (stripped) and the presence of 0.1 M KCl and 0.2 mM inositol hexaphosphate (+KCl +IHP). Stripped and +KCl +IHP treatments were prepared at three different pHs (for a total of 6 treatments per Hb sample), where working solutions was adjusted with NaOH to as near 7.2, 7.4, or 7.6 as possible, then pH was precisely measured with an Orion Star A211 pH Meter and Orion™ PerpHecT™ ROSS™ Combination pH Micro Electrode.

### Measuring O_2_-binding properties

O_2_-equilibrium curves were measured using a Blood Oxygen Binding System (Loligo Systems) at 37°C. The pH of whole-blood samples was set by measuring curves in the presence of 45 torr CO_2_, whereas the pH of Hb solutions was set with HEPES buffer (see above). Each whole-blood sample and Hb solution were sequentially equilibrated with an array of oxygen tensions (*P*_O2_) while the sample absorbance was continually monitored at 430 nm (deoxy peak) and 421 nm (oxy/deoxy isobestic point). Each equilibration step was considered complete when the absorbance at 430 nm had stabilized (2 - 4 minutes). Only oxygen tensions yielding 30 - 70% Hb–O_2_ saturation were used in subsequent analyses. Hill plots (log[fractional saturation/[1-fractional saturation]] vs. logP_O2_) were constructed from these measurements. A linear regression was fit to these plots and was used to determine the *P*_O2_ at half-saturation (*P*_50_) and the cooperativity coefficient (*n*_50_), where the X-intercept and slope of the regression line represent the *P*_50_ and *n*_50_, respectively. Whole-blood samples (n=3) are presented as mean±SE. For Hb solutions, a linear regression was fit to plots of log*P*_50_ vs. pH, and the resulting equation was used to estimate *P*_50_ values at pH 7.40 (± SE of the regression estimate).

## Acknowledgments

We would like to extend our sincere gratitude to the trainers and veterinary staff at SeaWorld of California for their help in this project, Jennifer Rego for blood sample collection and Dr. Judy St. Leger for logistical support. We thank S. Mohammadi, N. Gutierrez-Pinto, J. Hite, M. Kobiela, A. Dhawanjewar, M. Kulbaba, M. Gaudry and A. Quijada-Rodriguez for helpful comments on this manuscript. This research was supported by funding from the National Institutes of Health (HL087216 [JFS] and F32HL136202 [MST]), the National Science Foundation (OIA-1736249 [JFS], IOS-1927675 [JFS] and 1927616 [MST]) and the SeaWorld Parks & Entertainment technical contribution (2020-19).

