## Supplementary Information for "Evolved increases in hemoglobin-oxygen affinity and Bohr effect coincided with the aquatic specialization of penguins"

**This PDF file includes:**

Figures S1 to S4
Tables S1 to S3
SI References

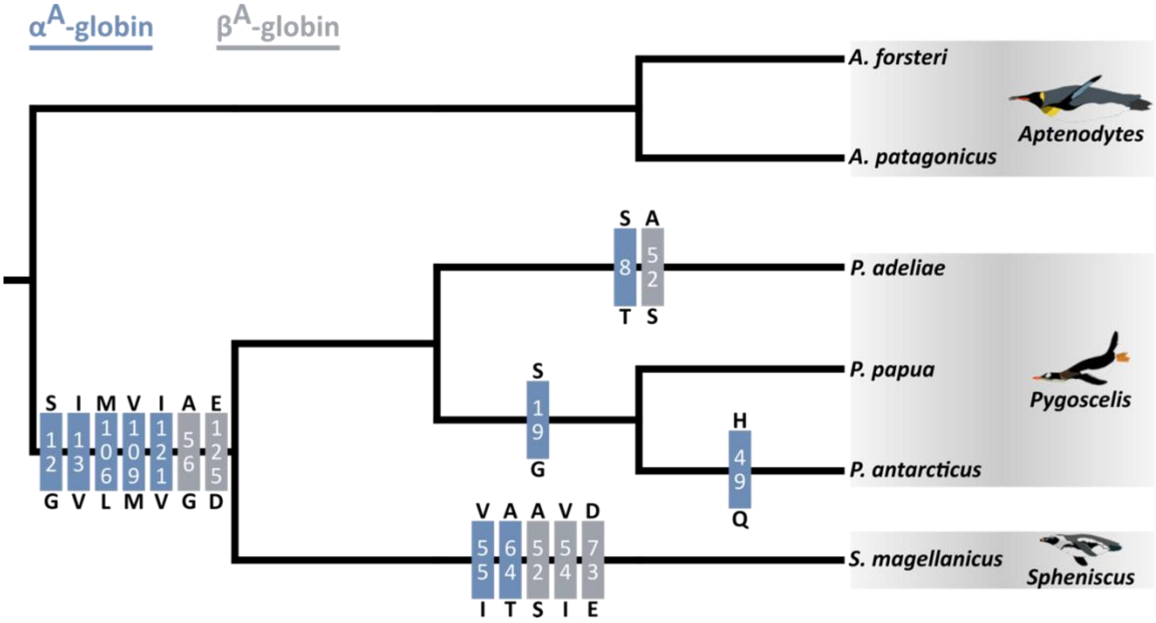


**Fig. S1.** Amino acid substitutions in the α- and β-globin chains of the six penguin species that were used in the experimental analysis of Hb function. Blue and grey numbered boxes represent amino acid positions in the α-globin and β-globin chains, respectively. Amino acids listed above and below each position represent the ancestral and derived amino acid state, respectively. The penguin phylogeny is adapted from Pan et al. (1).

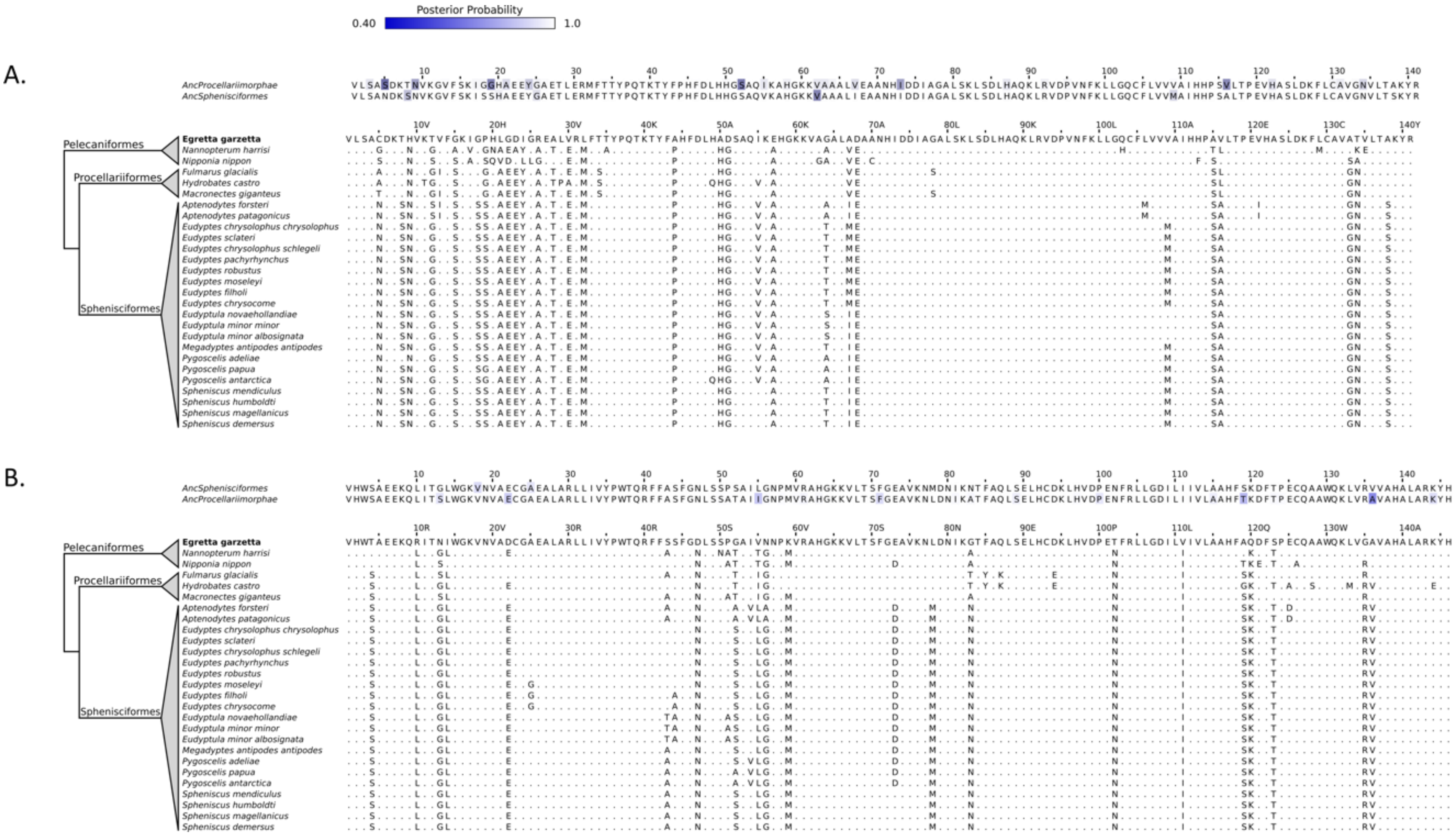

**Fig. S2.** Amino acid sequences for waterbird (A) α-globin and (B) β-globin chains. Estimated ancestral sequences for AncProcellariimorphae and AncSphenisciformes are shown above the native protein sequences. Shading on the ancestral protein sequences represent the posterior probability of each ancestral character estimate, where darker coloration represents lower posterior probability (see Figures S3 and S4 for more detail).

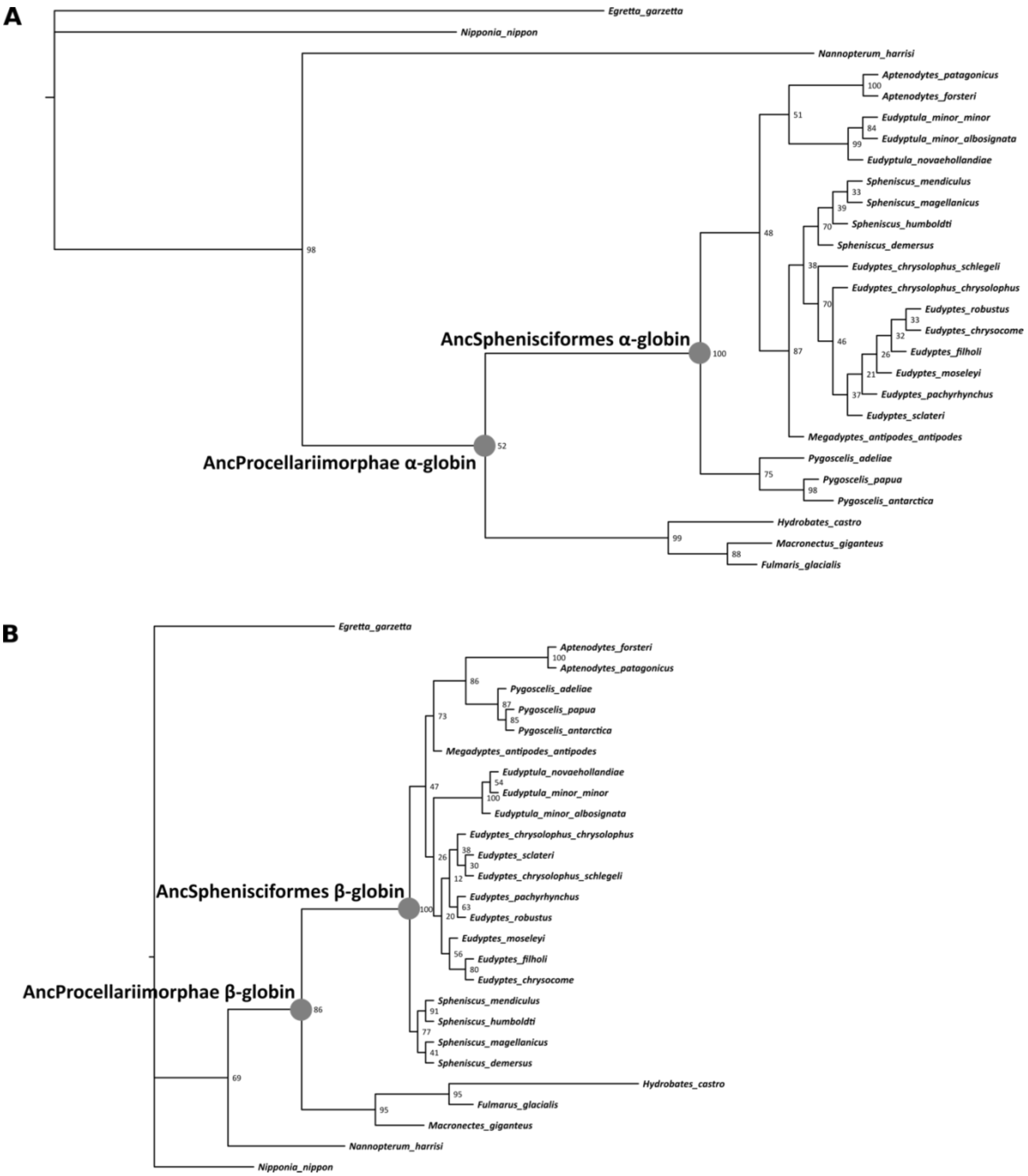


**Fig. S3.** Estimated maximum likelihood phylogenies of waterbird (A) $\alpha$-globin and (B) $\beta$-globin genes. Filled circles represent nodes for which ancestral sequences were reconstructed. Node labels represent percentage bootstrap support.

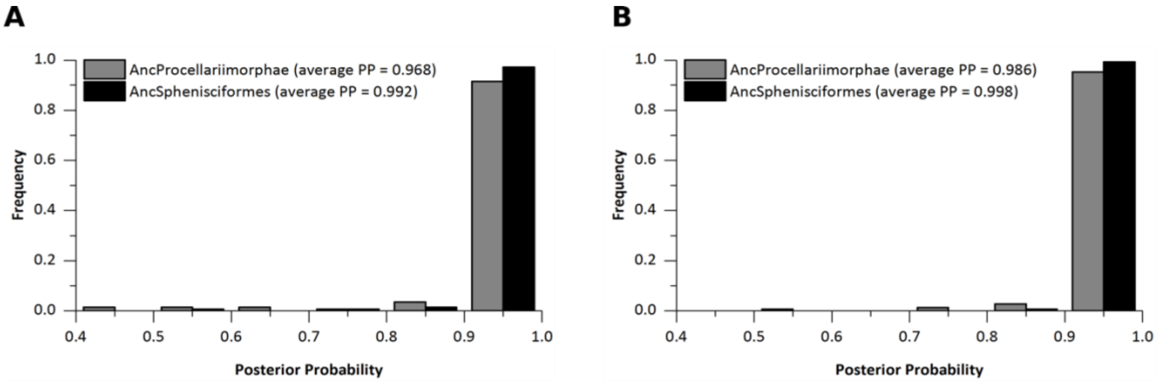


**Fig. S4.** Distributions of site-specific posterior probabilities for the (A) $\alpha$-globin and (B) $\beta$-globin genes of the estimated penguin ancestor (AncSphenisciformes) and the last common ancestor penguins shared with Procellariiformes (AncProcellariimorphae).

**Table S1.** Oxygen affinities ($P_{50}$, torr) of penguin whole-blood and native Hbs with their sensitivity to allosteric effectors at 37°C in 0.1 M HEPES buffer at pH 7.4.

| Species | [a]Whole-Blood $P_{50}$ | [b]Whole-blood $n_{50}$ | [a]Stripped $P_{50}$ | [a]Stripped $\log P_{50}$ | [b]Stripped $n_{50}$ | [c]Cofactor Effect | [d]Bohr Coefficient ($\Delta\log P_{50}/\Delta$pH) | |
|---|---|---|---|---|---|---|---|---|
| | | | | | | | Stripped | KCl + IHP |
| *A. forsteri* | 31.77±1.19 | 2.67±0.03 | 3.25±0.16 | 0.51±0.02 | 1.48±0.13 | 0.89 | -0.282±0.054 | -0.502±0.009 |
| *A. patagonicus* | 38.10±0.99 | 2.79±0.14 | 4.46±0.12 | 0.65±0.01 | 1.24±0.03 | 0.83 | -0.120±0.036 | -0.554±0.133 |
| *P. adeliae* | 33.16±3.73 | 2.37±0.11 | 3.50±0.36 | 0.54±0.04 | 1.44±0.12 | 0.82 | -0.300±0.124 | -0.625±0.037 |
| *p. papua* | 31.74±0.51 | 2.72±0.13 | 3.54±0.60 | 0.55±0.07 | 1.59±0.14 | 0.83 | -0.178±0.221 | -0.661±0.055 |
| *P. antarcticus* | 30.37±1.60 | 2.45±0.35 | 3.80±0.08 | 0.58±0.01 | 1.42±0.05 | 0.66 | -0.190±0.035 | -0.454±0.105 |
| *S. magellanicus* | 34.97±1.30 | 3.27±0.38 | 3.36±0.07 | 0.53±0.01 | 1.64±0.03 | 0.84 | -0.190±0.022 | -0.365±0.035 |

[a]$P_{O2}$ at half-saturation (torr)
[b]Hill's cooperativity coefficient
[c]$\Delta\log P_{50}$([KCl+IHP] – Stripped)
[d]Purified Hb, pH range 7.2 to 7.6

**Table 2**. Branch-site model statistics for reconstructed ancestral Sphenisciformes α- and β-globin genes. NS = Not significant.

| Model | np | -lnL | Estimates | LRT |
|---|---|---|---|---|
| α-globin Model A null | 55 | -1454.78 | Background: $\omega_0$=0.04, $\omega_1$=1.00, $\omega_{2a}$=0.04, $\omega_{2b}$=1.00<br>Foreground: $\omega_0$=0.04, $\omega_1$=1.00, $\omega_{2a}$=1.00, $\omega_{2b}$=1.00<br>$p_0$=0.63, $p_1$=0.29, $p_{2a}$=0.08, $p_{2b}$=0.03 | |
| α-globin Model A alt (ω1 > 1) | 56 | -1454.76 | Background: $\omega_0$=0.05, $\omega_1$=1.00, $\omega_{2a}$=0.04, $\omega_{2b}$=1.00<br>Foreground: $\omega_0$=0.05, $\omega_1$=1.00, $\omega_{2a}$=2.50, $\omega_{2b}$=2.50<br>$p_0$=0.68, $p_1$=0.26, $p_{2a}$=0.04, $p_{2b}$=0.02 | NS |
| β-globin Model A null | 55 | -1359.51 | Background: $\omega_0$=0.03, $\omega_1$=1.00, $\omega_{2a}$=0.03, $\omega_{2b}$=1.00<br>Foreground: $\omega_0$=0.03, $\omega_1$=1.00, $\omega_{2a}$=1.00, $\omega_{2b}$=1.00<br>$p_0$=0.85, $p_1$=0.15, $p_{2a}$=0.00, $p_{2b}$=0.00 | |
| β-globin Model A alt (ω1 > 1) | 56 | -1359.51 | Background: $\omega_0$=0.03, $\omega_1$=1.00, $\omega_{2a}$=0.03, $\omega_{2b}$=1.00<br>Foreground: $\omega_0$=0.03, $\omega_1$=1.00, $\omega_{2a}$=1.00, $\omega_{2b}$=1.00<br>$p_0$=0.85, $p_1$=0.15, $p_{2a}$=0.00, $p_{2b}$=0.00 | NS |

**Table 3**. Clade model statistics for reconstructed ancestral Sphenisciformes $\alpha$- and $\beta$-globin genes. NS = Not significant.

| Model | np | -lnL | Site Class | Proportion | $\omega$ | $\omega$ Branch Type 0 | $\omega$ Branch Type 1 | LRT |
|---|---|---|---|---|---|---|---|---|
| | | | 0 | 0.53 | 0.00 | | | |
| $\alpha$-globin Model 2a_rel | 56 | -1452.71 | 1 | 0.11 | 1.00 | | | |
| | | | 2 | 0.36 | 0.36 | | | |
| | | | 0 | 0.69 | | 0.04 | 0.04 | |
| $\alpha$-globin Model C | 57 | -1451.82 | 1 | 0.15 | | 1.00 | 1.00 | NS |
| | | | 2 | 0.17 | | 1.01 | 0.00 | |
| | | | 0 | 0.65 | 0.03 | | | |
| b-globin Model 2a_rel | 56 | -1359.51 | 1 | 0.15 | 1.00 | | | |
| | | | 2 | 0.2 | 0.03 | | | |
| | | | 0 | 0.43 | | 0.03 | 0.03 | |
| $\beta$-globin Model C | 57 | -1358.49 | 1 | 0.15 | | 1.00 | 1.00 | NS |
| | | | 2 | 0.42 | | 0.06 | 0.00 | |
